# Sphingolipids regulate the tethering stage of vacuole fusion by affecting membrane fluidity

**DOI:** 10.1101/2020.02.17.953331

**Authors:** Chi Zhang, Logan R. Hurst, Zeynep D. Gokbayrak, Jorge D. Calderin, Michael R. Hrabak, Adam Balutowski, David A. Rivera-Kohr, Thomas D.D. Kazmirchuk, Christopher L. Brett, Rutilio A. Fratti

## Abstract

Sphingolipids are essential in membrane trafficking and cellular homeostasis. Here, we show that sphingolipids containing very long-chain fatty acids (VLCFAs) promote robust homotypic vacuolar fusion in *Saccharomyces cerevisiae*. The elongase Elo3 adds two carbons to 24-carbon (C24) acyl chains to make C26 VLCFAs that are incorporated into sphingolipids. Vacuoles isolated from *elo3*Δ cells had increased fluidity relative to the wild-type and were attenuated for fusion. Upon further testing we found that vesicle the tethering stage was affected as *elo3*Δ vacuole clusters contained fewer vesicles versus the WT. Vacuole tethering requires the interactions of late endosomal Rab GTPase Ypt7 and the HOPS tethering complex. Pulldown assays using GST-Ypt7 showed that HOPS from *elo3*Δ vacuole extracts failed to bind Ypt7 while HOPS from WT extracts interacted with GST-Ypt7. Furthermore GFP-Ypt7 failed to localize at vertex microdomains of *elo3*Δ vacuoles relative to the WT, whereas HOPS and regulatory lipids did accumulate at vertices. Finally, we found that *elo3*Δ vacuoles had reduced V-ATPase. Together these data show that C26-VLCFA containing sphingolipids are important for Ytp7 function and vacuole homeostasis.

## Introduction

The lipid composition of cell membranes is extremely complex, and provides the physical properties needed by each membrane so it can perform is functions (Harayama & Riezman, 2018). Tightly controlling the stoichiometry of lipids is a key determinant in the ability of a cell to carry out trafficking events such as exocytosis and organelle fusion, and slight changes to the ratio of lipids can drastically affect these essential processes (Johansen *et al*, 2012; Starr & Fratti, 2019a). Using the budding yeast *Saccharomyces cerevisiae* as a model system facilitates the study of how select classes and species of lipids are individually modulated, and how the effects of these adjustments can be monitored at the level of specific organelles, such as the eukaryotic lysosome. The lysosome is often viewed as a catabolic endpoint for endocytic and autophagic events, but it also serves as a hub for many cellular signaling processes, including cellular ageing and longevity, as well as pathogen clearance through the phagosome-lysosome fusion pathway (Carmona-Gutierrez *et al*, 2016; Li & Kane, 2009; Uribe-Querol & Rosales, 2017). Sphingolipids have been identified as bioactive lipids in mammalian cells, with roughly 40 enzymes contributing to their highly regulated metabolism, and defects in their metabolism is connected to many human lysosomal storage diseases. That said, the correlation between sphingolipids and lysosome morphology and function is not well understood in molecular detail (Hannun & Obeid, 2018; Platt, 2014).

Membrane fusion between organelles/vesicles occurs by progressing through a series of experimentally defined stages that are conserved in the trafficking of secretory, endosomal and autophagic membranes (Conradt *et al*, 1994; Starr *et al*, 2019, 2016; Wickner, 2010; Wickner & Schekman, 2008). In the case of *S. cerevisiae*, the spatiotemporal localization and activity of many proteins and lipids that participate in homotypic vacuole fusion are well understood. Prepriming of the membranes is dependent on the conversion of phosphatidic acid (PA) to diacylglycerol (DAG) by the PA phosphatase Pah1, which allows the transfer of the AAA+ ATPase Sec18 (NSF) from the membrane to inactive *cis*-SNARE complexes via its adaptor protein Sec17 (α-SNAP) (Sasser *et al*, 2012a; Starr *et al*, 2019, 2016). Priming occurs when Sec18 hydrolyzes ATP and transfers mechanical force through Sec17 onto the cis-SNARE bundle to separate it into its individual constitutive SNAREs (Mayer *et al*, 1996; Mayer & Wickner, 1997). This is followed by tethering of the vacuoles through interactions between the late endosomal/lysosomal Rab GTPase Ypt7/Rab7 and its heterohexameric HOPS effector complex (Hickey *et al*, 2009; Price *et al*, 2000a; Seals *et al*, 2000; Stroupe *et al*, 2006; Wurmser *et al*, 2000). During docking vacuoles are drawn together to form flattened discs termed boundary domains where they are in tight contact. At the edge of the boundary disc, where vacuoles come together are membrane microdomains that form during this phase of fusion and become enriched in the lipids and proteins that drive membrane fusion (Fratti *et al*, 2004; Jun *et al*, 2006; Karunakaran *et al*, 2012; Wang *et al*, 2003b, 2002b). During this, SNARE proteins from opposite membranes bundle together in a tight, coiled-coil motif called a *trans*-SNARE complex that triggers the release of Ca^2+^ from the vacuole lumen (Collins & Wickner, 2007; Merz & Wickner, 2004; Ungermann *et al*, 1998). Hemifusion occurs when the outer cytoplasmic facing leaflets fuse while leaving the inner leaflets intact (Mattie *et al*, 2017; Reese *et al*, 2005; Reese & Mayer, 2005). This ultimately leads to pore formation between the two organelles and full content mixing to complete a fusion cycle. Tethering, docking, and fusion steps also require the activity of the Rho-family GTPases Cdc42 and Rho1, which control cycles of actin remodeling at the vacuole membrane (Eitzen *et al*, 2002; Isgandarova *et al*, 2007; Logan *et al*, 2010).

In addition to proteins, membrane fusion requires a set of regulatory lipids, which are often enzymatically modified by kinases, phosphatase and lipases at specific times throughout the endosomal maturation and homotypic vacuole fusion processes (Starr & Fratti, 2019b). These lipids co-enrich with fusion machinery proteins in an interdependent manner to form the “vertex domain”, the ring-like area surrounding two membranes in contact during the docking stage (Fratti *et al*, 2004; Wang *et al*, 2002b). The regulatory glycerophospholipids DAG, PA, and a number of phosphatidylinositol phosphates (PI) have been shown to influence the recruitment and activity of Ypt7, its nucleotide exchange factor Mon1-Ccz1 and effector complex HOPS, the SNARE chaperone proteins Sec18 and Sec17, the soluble SNARE Vam7, and actin (Boeddinghaus *et al*, 2002; Cabrera *et al*, 2014; Cheever *et al*, 2001; Fratti *et al*, 2004; Jun *et al*, 2004; Karunakaran *et al*, 2012; Karunakaran & Fratti, 2013; Lawrence *et al*, 2014; Li *et al*, 2014; Miner *et al*, 2019, 2016; Starr & Fratti, 2019b; Stroupe *et al*, 2006). In contrast to glycerophospholipids, there are only a few studies that investigate the role of sphingolipids in the system of homotypic vacuole fusion. In wild type yeast very-long-chain fatty acids (VLCFAs) are almost exclusively bound to long chain bases (LCBs) in the form of ceramides or glycosphingolipids. Free VLCFAs in cell extracts are scarcely detectable (Lester *et al*, 1993). Baker’s yeast contains three fatty acid elongases (Elo1, Elo2, and Elo3), which catalyze the condensation of a malonyl-CoA unit with a long-chain fatty acyl-CoA at the endoplasmic reticulum. Each of these enzymes has an inherent substrate specificity and unique VLCFA major product, such as Elo3 displaying specificity for long- to very-long-chain fatty acid substrates and producing VLCFAs up to 26-carbons in length (Denic & Weissman, 2007; Rössler *et al*, 2003). In mammalian cells VLCFA production has been linked to processes including phagocytosis, apoptosis, and myelin function, and has been connected to diseases such as ichthyosis, macular degeneration, and myopathy (Kihara, 2016; Sassa & Kihara, 2014; Tafesse *et al*, 2015). The C26 VLCFA is likely critical for the function of yeast sphingolipids in membranes based on research demonstrating that this particular species of fatty acid is found in the vast majority sphingolipid species in wild-type yeast (Ejsing *et al*, 2009). Studies with yeast lacking functional elongase enzymes have uncovered links between sphingolipids and Vps21-related endosomal maturation, vacuolar acidification, and vacuolar morphology/function, but their role as regulatory lipids in membrane fusion remains unclear (Chung *et al*, 2003; Faber *et al*, 2009; Gaigg *et al*, 2001; Kohlwein *et al*, 2001; Obara *et al*, 2013).

Giant unilamellar vesicles (GUVs), membranes/lipids isolated from cells, and vacuoles in live yeast have been found to form lipid rafts, also known as membrane microdomains and/or nanodomains. These are transient, sub-organellar domains, which phase separate within a membrane based on differences in lipid and protein composition and cause differences in properties such as fluidity and thickness. These domains are highly enriched in sterols, sphingolipids and other lipids with saturated acyl chains, and display liquid-ordered (L_o_)-like properties (Klose *et al*, 2010; Sezgin *et al*, 2017; Toulmay & Prinz, 2013). Thus, it is likely that sphingolipids colocalize with ergosterol in the vertex ring to regulate fusion. It is also likely that vacuolar sphingolipids affect other vacuolar functions including acidification by the V-ATPase (Chung *et al*, 2003; Finnigan *et al*, 2012), Ca^2+^ transport (Deng *et al*, 2018; Shatrov *et al*, 1997) and actin remodeling (Balguerie *et al*, 2002; Fu *et al*, 2018).

In this study we present evidence that a lack of C26 VLCFA-containing sphingolipids in *elo3*Δ yeast affects membrane vacuole fluidity, morphology and homotypic fusion. Inhibited fusion was largely due to defects in the tethering stage of vacuole fusion, which is linked Ypt7 function. This is demonstrated by reduced vacuole clustering, defective interactions between Ypt7 and HOPS, and the lack of Ytp7 enrichment at vertex sites. Together our data indicate that C26 VLCFA containing sphingolipids affect the assembly specialized membrane microdomains that are necessary in regulating the machinery that drives vacuole fusion.

## Results

### Yeast cells lacking Elo3 are enlarged and have misshapen vacuoles

To probe the role that C26 VLCFA-containing sphingolipids (C26-SL) play in vacuole fusion the *ELO3* gene (formerly *SUR4*) was deleted from *S. cerevisiae,* and vacuole morphology was examined in live cells. Wild type (WT) BJ3505 yeast grown to logarithmic phase in YPD were round, and most cells had one or two vacuoles that occupied roughly half of the cell volume as visualized by fluorescent microscopy with the lipophilic dye FM4-64 (Vida & Emr, 1995) (**Fig. 1A-C**). However, BJ3505 *elo3*Δ cells were much larger, and were often with adjacent to smaller vesicles, which corresponded to a Class F vacuole morphology phenotype (Raymond *et al*, 1992) (**Fig. 1A-C**). Others have described the shape of *ELO3* mutants as “dumbbell-shaped” (Desfarges *et al*, 1993). We also saw this in some elongated cells (data not shown); however, most cells harbored the Class F phenotype.

**Figure 1.**
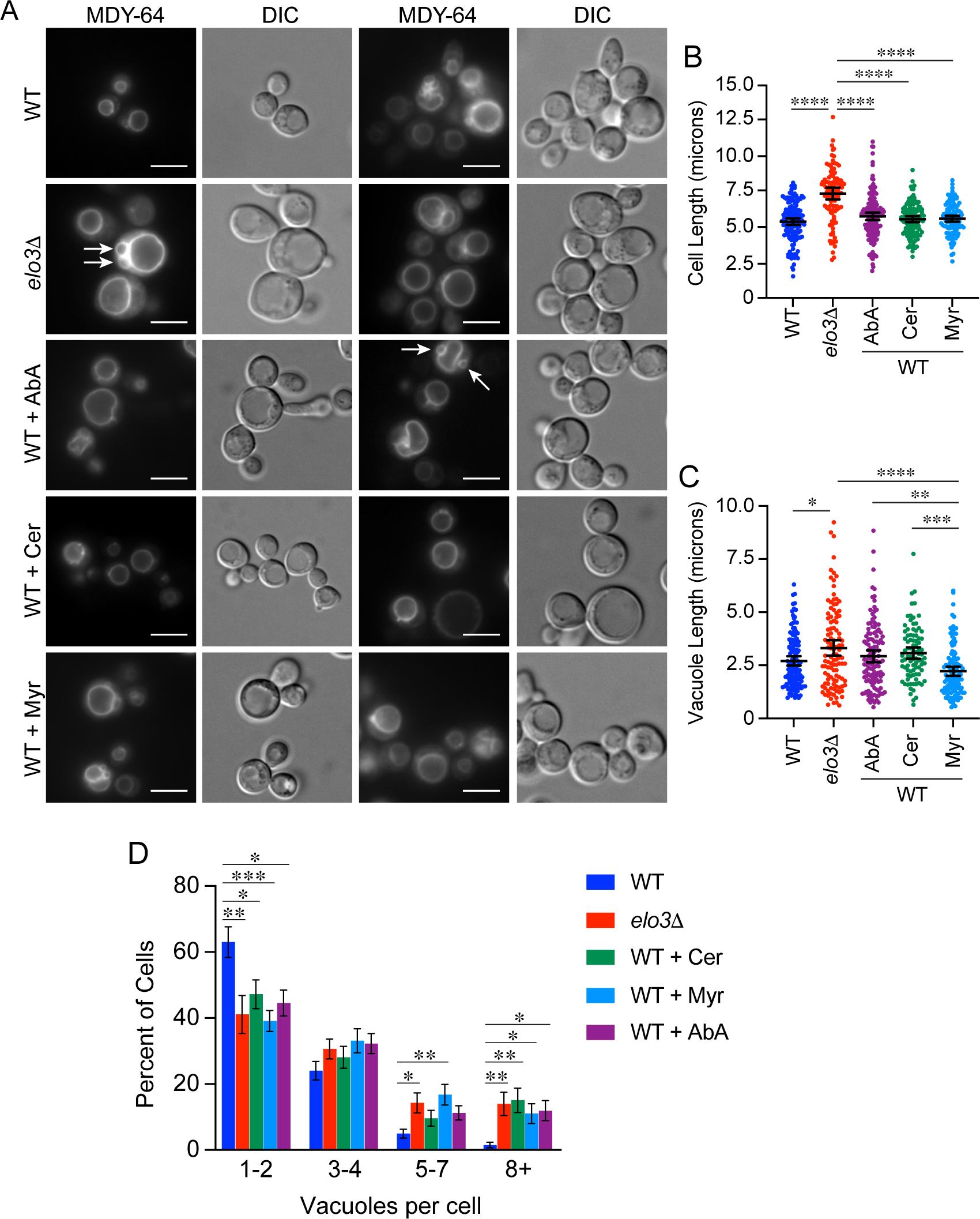
Cellular and vacuole morphology are perturbed when sphingolipid synthesis is interrupted. **(A)** BJ3505 wild type (WT) and *elo3*Δ yeast grown in YPD medium. Cells were grown overnight, back-diluted to OD600 ∼0.7 in fresh medium and grown for an additional 2 h. WT cells were then treated with 0.125 μg/mL aureobasidin A (AbA), 0.25 μg/mL myriocin (Myr), 50 µM cerulenin (Cer) or methanol (vehicle) and incubated for an additional 4-5 h. Cells were then incubated with 10 µM MDY-64 for 15 min in the dark before visualization. Images are representative of three separate trials. Arrows indicate the Class F phenotype vacuoles. Scale bars: 4 μm. **(B-C)** Cell and vacuole size were measured using images from **(A)**. Measurements were plotted as a scatter plot error bars representing the mean ± 95% CI. **(B)** Vacuole length: One-way ANOVA for multiple comparisons with no treatment (0 µM) as a control [F (4, 558) = 9.094; \*\*\*\**p*<0.0001]. Tukey’s multiple comparison test was used for individual p values n = ∼150 cells per condition. \**p*<0.05, \*\**p*<0.01, \*\*\*\**p*<0.001. **(C)** Cell length: One-way ANOVA for multiple comparisons [F (4, 630) = 31.72; \*\*\*\**p*<0.0001]. Tukey’s multiple comparison test was used for individual *p* values n = ∼150 cells per condition. \*\*\*\**p*<0.001. **(D)** Vacuole fragmentation was measured using images from **(A)**. Measurements were plotted as bars representing mean ± SEM. Significance was determined with one-way ANOVA for multiple comparisons [F (4, 130) = 3.749, ** *p*=0.0064]. Error bars represent SEM. (n=3). Dunnett’s multiple comparison test was performed for each cluster bin for individual *p* values n= ∼150 cells per condition.

We next visualized live WT yeast after treatment with known inhibitors of *de novo* sphingolipid biosynthesis. Myriocin was used to inhibit the rate-limiting step of LCB synthesis by the serine palmitoyltransferase complex (Lcb1/Lcb2) (Kvam *et al*, 2005; Nagiec *et al*, 1994). Aureobasidin A (AbA) was used to block inositol-phosphoceramide (IPC) synthesis by the Aur1/Kei1 IPC synthase complex (Hashida-Okado *et al*, 1996; Voynova *et al*, 2015). Finally, we used cerulenin, which has been reported to inhibit both the soluble fatty acid synthase (FAS) complex and the microsomal VLFCA elongase enzymes at high concentrations (Kvam *et al*, 2005). While all three inhibitors perturbed cell and vacuole morphology the differences were not statistically significant. Treatment with myriocin caused a vacuole fragmentation phenotype (Class B vacuole morphology) and very small cells. This was intriguing when we considered that IPC has been shown to be enriched at the yeast vacuole (Hechtberger *et al*, 1994), and IPC and ceramide have been suggested to be required for proper vacuole morphology and function (Faergeman *et al*, 2004). When we quantitated the effects of these drugs on the number of vacuoles per cell, we found that they all reduced the number of cells with only one vacuole to match *elo3*Δ cells (**Fig. 1D**). This was accompanied by significant increases in the number of cells that contained 5 or more vacuoles.

### Lipid packing and membrane fluidity is altered in the absence of C26-SL

Lipid rafts at the plasma membrane of eukaryotes are characterized by their increased rigidity and thickness in relation to its surrounding membrane environment (Simons & Vaz, 2004). These characteristics are due in part to tight packing of sphingolipids, sphingomyelin in particular, with cholesterol. Although not identical to classically defined lipid rafts, yeast vacuoles form ergosterol-rich microdomains at the vertex contact points of docked vacuoles (Fratti *et al*, 2004; Wang *et al*, 2003b, 2002b). Here we tested whether the absence of C26 VLCFAs affected vacuolar lipid packing and fluidity. To measure membrane changes in lipid packing and fluidity, we used merocyanine 540 (MC540) (Langner & Hui, 1999; Williamson *et al*, 1983; Wilson-Ashworth *et al*, 2006; Stillwell *et al*, 1993). MC540 fluoresces at two major peaks that correlate with liquid crystalline and gel-phase membranes. MC540 is excluded from membranes with very tight lipid packing and has a peak emission at 624 nm when inserted in gel-phase membranes. MC540 fluorescence shifts to 594 nm when it intercalates in liquid crystalline membranes with increased fluidity linked to looser lipid packing. Thus, increased fluidity would correspond with an increase in fluorescence at 594 nm. We chose MC540 because its fluorescence was not affected by small molecule inhibitors that we commonly use including dibucaine, an anesthetic that is known to increase membrane fluidity (Papahadjopoulos *et al*, 1975). The fluorescence of other probes used to measure fluidity, e.g., pyrenedecanoic acid, and TMA-DPH (1-(4-trimethyammoniumphenyl)-6-phenyl-1,3,5-hexatriene p-toluenesulfonate) overlap with that of dibucaine making it difficult to use in these experiments.

In Figure 2 we show MC540 fluorescence at 594 nm with WT and *elo3*Δ vacuoles. We found that the fluorescence of MC540-labeled *elo3*Δ vacuoles at 594 nm was significantly higher relative to the WT signal indicating that the lack of C26 VLCFAs in *elo3*Δ vacuoles made them more fluid (**Fig. 2A**). This was consistent with the role of SL at the plasma membrane and the formation of lipid raft-like domains. To verify that we were measuring differences in fluidity, we tested vacuoles in the presence of dibucaine. We found that dibucaine increased the fluorescence of MC540-labeled WT vacuoles suggesting that the anesthetic significantly increased membrane fluidity (**Fig. 2B**). When *elo3*Δ vacuoles were treated with dibucaine we found that there was no further increase in fluidity.

**Figure 2.**
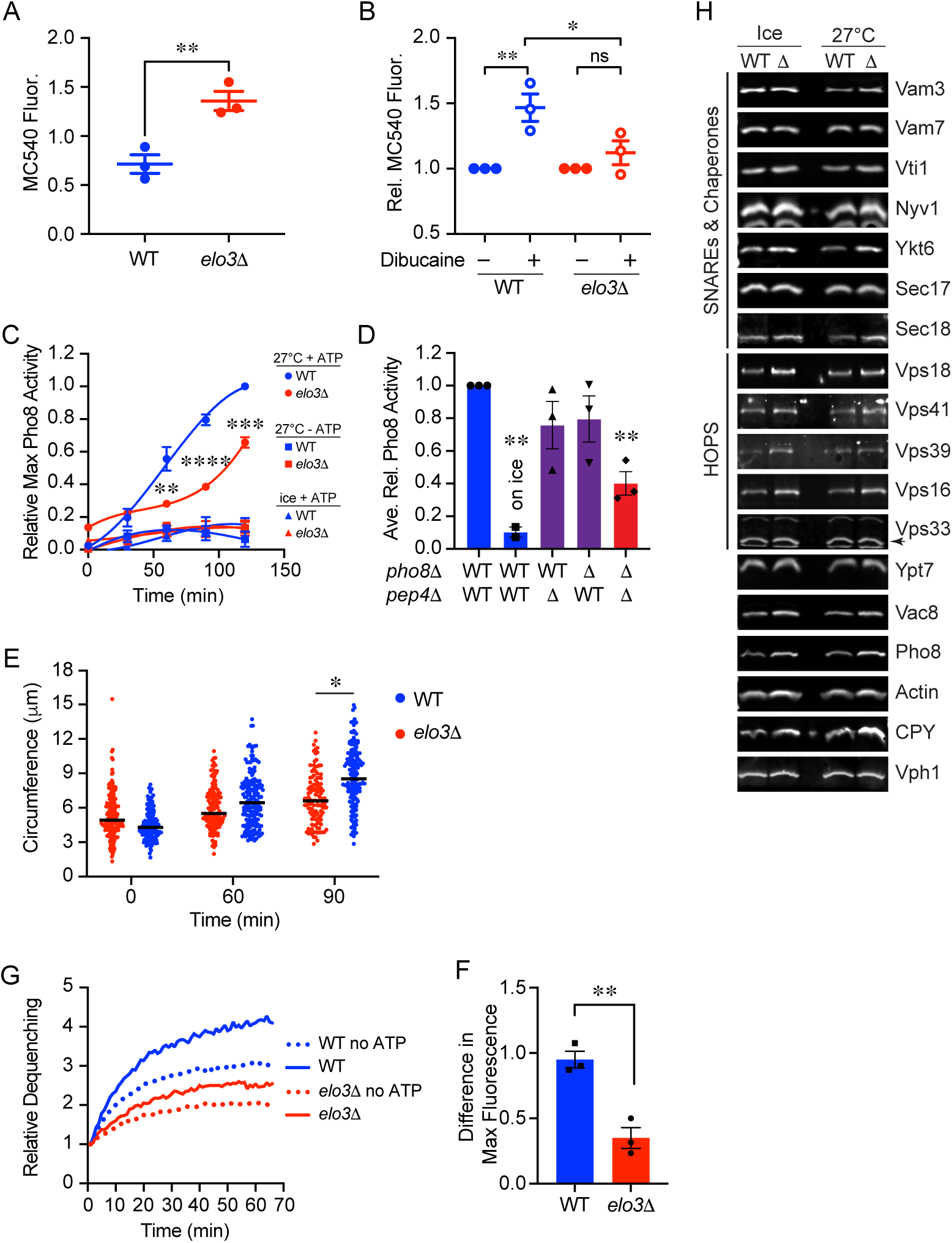
Vacuoles from *elo3*Δ cells have increased fluidity and are fusion impaired. **(A)** Vacuoles from *elo3*Δ cells have increased fluidity relative to WT. Isolated vacuoles were mixed with MC540 in PS buffer and incubation in the dark for 20 min at 25°C. Vacuoles were reisolated and resuspended in PS buffer with ATP, MgCl2 and 10X fusion, CoA. MC540 fluorescence was measured with an excitation of 530 nm and emissions at 585. WT and elo3Δ were solubilized 1% TX100 after labeling to release the incorporated dye in each set of vacuoles. Fluorescence intensities were plotted as values relative to maximum fluorescence in detergent. Error bars represent SEM (n=3). Data was analyzed using an unpaired two-tailed t-test. \*\**p*<0.01. **(B)** WT vacuole fluidity was determined as described above in the presence or absence of 500 µM dibucaine in 0.5% ethanol. Here, changes in fluorescence were normalized to each strain that was set to 1. Changes in fluorescence was analyzed using one-way ANOVA for multiple comparisons with no treatment (0 µM dibucaine/0.5% ethanol) as a control [F (4, 14) = 14.87; \*\*\*\**p*<0.0001]. Error bars are mean ± SEM. Dunnett’s multiple comparison test was used for individual *p* values (n=3). \*\**p*<0.01. **(C)** A time course of fusion activity was measured by mixing 3 μg of DKY6281 vacuoles and 3 μg of BJ3505 vacuoles or their *elo3*Δ derivatives then incubating samples at 27°C until the indicated time point, when the reactions were placed on ice. Fusion activity was expressed as the percentage of WT fusion activity by measuring the absorbance at 400 nm for each sample after 120 min incubation at 27°C. Significance was determined using one-way ANOVA for multiple comparisons for each time point between strains. Error bars represent SEM (n=3). [F (5, 12) = 15.62; \*\*\*\**p*<0.0001]. Error bars are mean ± SEM. Tukey’s multiple comparison test was used for individual *p* values (n=3). \*\**p*<0.01. \*\*\**p*<0.001, \*\*\*\**p*<0.0001 **(D)** Endpoint fusion reactions were performed with combinations of wild-type (WT) and *elo3*Δ (11) vacuoles from the indicated strains at the bottom of the panel. Significance was determined using one-way ANOVA for multiple comparisons with WT/WT as a control. Error bars represent SEM (n = 3). [F (4, 9) = 10.07; **p=0.0022]. Error bars represent mean ± SEM. Dunnett’s test for multiple comparison was used for individual *p* values. \*\**p*<0.01. **(E)** Measurement of vacuole diameters after fusion. Vacuoles were incubated for the indicated times at 27°C. Diameters of individual vacuoles in clusters were measured for WT and *elo3*Δ. Each measurement was illustrated in a scatter plot. The bars represent the mean. (n=3). \**p*<0.05. **(F)** Lipid mixing experiments were performed with vacuoles isolated from WT BJ3505 or *elo3*Δ BJ3505 yeast. Reporter vacuoles (2 μg) of the indicated strain labeled with 150 μM Rh-PE were incubated with 16 μg unlabeled vacuoles from the same background. Fluorescence (λ_ex_=544 nm, λ_em_=590 nm) was measured every 60s for 40 min following the addition of ATP (at t = 5 min). **(G)** Quantitation of three experiments in **(F)** (n = 3). Error bars represent mean ± SEM. Data was analyzed using an unpaired two-tailed t-test \*\**p*<0.01. **(H)** Western blots of vacuole proteins using WT and *elo3*Δ vacuoles that were incubated on ice or at 27°C for 1h. Western blot with antibodies raised against the indicated protein. Rabbit antibodies against Vam3, Vam7, Vti1, Ykt6, Sec17, Sec18, Vps18, Vps41, Vps39, Vps16, Vps33, Ypt7, Vac8, Pho8 and Actin were prepared as previously described (Haas *et al*, 1995; Mayer *et al*, 1996; Mayer & Wickner, 1997; Nichols *et al*, 1997; Price *et al*, 2000b; Seals *et al*, 2000; Ungermann *et al*, 1999; Ungermann & Wickner, 1998; von Mollard *et al*, 1997; Wang *et al*, 2003a, 2002a, 1998). Mouse Anti-Vph1 was from Abcam (ab113683), and Mouse anti-CPY (Carboxy-peptidase Y) was from Thermo-Fisher (A-6428). Goat anti-rabbit IgG (H+L) secondary antibody DyLight 650 (84546) conjugate or Goat anti-mouse IgG (H+L) secondary antibody DyLight 650 (84545) conjugate were from Thermo-Fisher. Fluorescence was measured with a BioRad Chemidoc MP or Azure 400.

### Vacuoles from *elo3*Δ cells have attenuated fusion activity

To examine how the lack of functional Elo3 affected homotypic fusion, vacuoles were isolated from WT and *elo3*Δ strains in BJ3505 (*PHO8 pep4*Δ) and DKY6281 (*pho8Δ PEP4*) backgrounds as described (Conradt *et al*, 1992; Haas *et al*, 1994). Upon fusion, the Pep4 protease in DYK8281 vacuoles will cleave the alkaline phosphatase zymogen pro-Pho8 to produce active Pho8. Isolated vacuoles were incubated under standard fusion conditions, and *in vitro* fusion was monitored via Pho8 phosphatase activity upon content mixing. Using this method, we found that *elo3*Δ vacuole fusion could only reach ∼50% relative to WT vacuoles by 90 min (**Fig. 2C**). The fusion of *elo3*Δ seemed to accelerate, or catch up after 120 min, however, overall fusion was still reduced by ∼30% compared to WT fusion.

We next asked if changes in membrane composition needed to be on both sets of vacuoles to inhibit fusion. To do this we mixed fusion partners from WT and *elo3*Δ backgrounds. Fusion reactions between vacuole sets that were isolated solely from the *elo3*Δ background were reduction in fusion relative to WT (**Fig. 2D, lane 5)**. However, when fusion partners were mixed from WT and *elo3*Δ backgrounds the fusion defect was mostly rescued (**Fig. 2D, lanes 3 & 4)**. Therefore, the Elo3-dependent factor(s) important for fusion can either be shared between vacuoles or sufficient on one side in supporting fusion. Together this data suggested that C26-SL were positive regulators of homotypic vacuolar fusion. To further verify that deleting *ELO3* negatively impacted vacuole fusion, we took visual measurements of vacuole circumferences at different time points during fusion as described (Miner *et al*, 2017; Sasser *et al*, 2012b). At each time point vacuoles were placed on ice and stained with FM4-64 prior to image acquisition by fluorescence microscopy. At the start of fusion (T=0 min) we observed that there was no significant difference in circumference between WT and *elo3*Δ vacuoles (**Fig. 2E**). As fusion reactions progressed, we measured a significant increase of WT vacuole circumference versus the deletion.

The loss of C26-VLCFAs increased the fluidity of *elo3*Δ vacuoles, which could affect multiple aspects of the fusion pathway. One phase that could be altered is the energy barrier at the hemifusion stage. Previously we showed that mutations that impaired SNARE function had no effect on hemifusion, yet they blocked full content mixing (Karunakaran & Fratti, 2013). To rescue these mutants, we used the cationic amphiphile chlorpromazine to alter membrane curvature and fluidity. That study showed that membrane curvature and fluidity affected fusion in part by setting the energy barrier that SNARE pairing needed to overcome to trigger the transition between hemifusion and full content mixing. Here we next tested if the lack of C26-SL affected fusion before or after the hemifusion intermediate. To do this we isolated vacuoles from WT BJ3505 and BJ3505 *elo3*Δ and labeled them with rhodamine B-conjugated phosphatidylethanolamine (Rh-PE) at a self-quenching concentration (Reese *et al*, 2005; Reese & Mayer, 2005; D’Agostino & Mayer, 2019). Labeled vacuoles were then incubated with an 8-fold excess of unlabeled vacuoles. As the outer leaflets fuse, Rh-PE is diluted leading to its dequenching and increased fluorescence. This is kinetically separated from full content mixing to show that hemifusion occurred (Jun & Wickner, 2007; Mattie *et al*, 2017). Using this method, we found that Rh-PE-labeled *elo3*Δ vacuoles were impaired in hemifusion compared to WT vacuoles (**Fig. 2F-G**). The VLCFAs present in sphingolipids have been suggested to help stabilize highly curved membranes, which promotes membrane fusion (Golani & Schwarz, 2023; Molino *et al*, 2014), which may help explain their contribution to fusion events. From these data we deduced that 26C-SLs regulated a stage upstream of hemifusion.

Because defects in vacuole fusion can be due to defective sorting of proteins to the vacuole through the AP3 or CPY pathways, we next checked the composition of *elo3*Δ vacuoles by Western blotting. We used freshly isolated WT and *elo3*Δ vacuoles and compared them to vacuoles that had been incubated for 1h at 27°C under fusion conditions. This was to see if there were any changes in degradation or protein modification. As seen in **Figure 2H** we did not observe any significant difference between WT and *elo3*Δ vacuoles under either condition. This told us that differences in fusion were not due to the absence of a fusion-driving protein.

### Vacuole acidification is reduced in *elo3*Δ vacuoles

We next investigated SL-dependent cellular functions that are associated/required for vacuole fusion including V-ATPase activity. The V-ATPase is composed of the integral membrane complex V_O_ and the soluble V_1_ complex (Vasanthakumar & Rubinstein, 2020). Investigators found that deleing *ELO3/SUR4* in W303-1A yeast led to the inability to acidify vacuoles as measured by quinacrine staining of whole cells (Chung *et al*, 2003). This was due to defective assembly of the V_1_ soluble head group. Isolated *elo3*Δ vacuoles lacked the Vma1, Vma2 and Vma5 V_1_ subunits, whereas WT vacuoles contained the V_1_ subunits. In conclusion, they found that the V_1_ complex was recruited to V_1_ complexes in a C26 VLCFA dependent manner. Others have found that regulating sphingolipid homeostasis is needed for full V-ATPase activity (Finnigan *et al*, 2011). Here we tested if isolated vacuoles could become acidified using an acridine orange fluorescence shift assay (Müller *et al*, 2002; Zhang *et al*, 2022b, 2022a). Acridine orange becomes trapped in acidic compartments due to its protonation. Monomeric acridine orange fluoresces at 520 nm whereas protonated acridine orange forms dimers and fluoresces at 680 nm. Thus, a decrease in fluorescence at 520 nm serves as a reporter of vesicle acidification. In Figure 3A we show that after the addition of ATP to the reaction, the fluorescence signal of the acridine orange decreased as protons entered the lumen. Using this assay, we found that vacuoles isolated from the *elo3*Δ strain have significantly impaired proton-pumping ability compared to WT vacuoles (**Fig. 3A, B**). This is consistent with the positive role of SL content in V-ATPase function and vacuole acidification.

**Figure 3.**
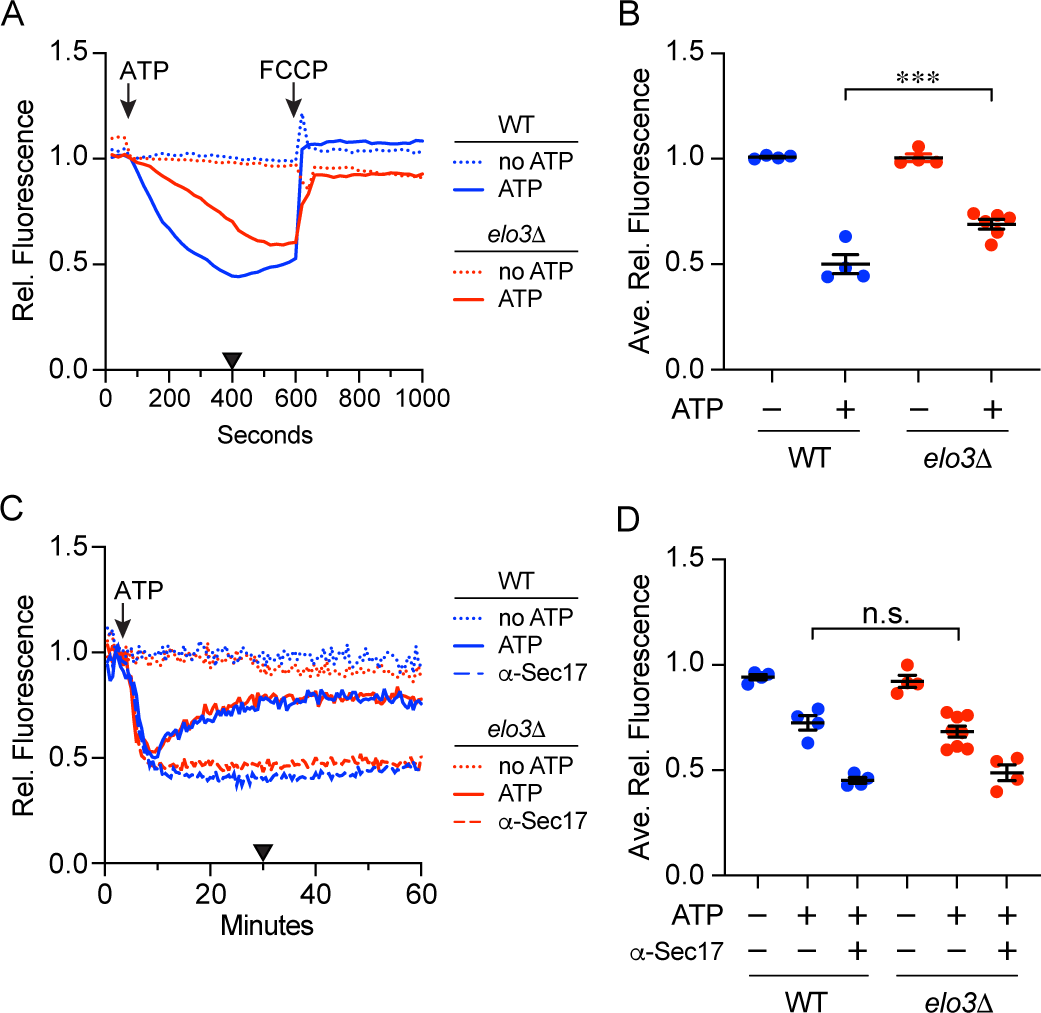
C26-SL are needed for optimal vacuole acidification by not Ca^2+^ transport. **(A)** Vacuole acidification was determined for WT and *elo3*Δ vacuoles by measuring changes in Acridine Orange fluorescence at 520 nm. Reactions were incubated in the presence or absence of ATP regenerating system and fluorescence was measured every 20 s over a 1000 s experiment. The protonophore FCCP was added at 500 s to collapse the proton gradient. Fluorescence was normalized to initial intensities at the time of adding ATP and set to 1. **(B)** Quantitation of multiple experiments shown in (A). Fluorescence values at 400 s were averaged for each strain and showed a significant difference between WT and mutant vacuoles. Significance was measured using one-way ANOVA for multiple comparisons [F (3, 14) = 80.30, **** *p*<0.0001]. Tukey’s post hoc multiple comparisons test was used for individual *p* values. Error bars represent mean SEM. (n > 4). \*\*\**p*<0.001 **(C)** Ca^2+^ transport across WT and *elo3*Δ vacuole membranes. Cal520 fluorescence was used to measure Ca^2+^ outside of vacuoles. Cal-520 fluorescence was measured in the absence of ATP, with ATP regenerating system, or anti-Sec17 and ATP regenerating system together. The lack of ATP prevented Ca^2+^ from entering the vacuole and generated a flat line with no change in fluorescence. As Ca^2+^ was transported into the vacuole lumen, Cal520 fluorescence was diminished. Subsequent Ca^2+^ release resulted in increased Cal520 fluorescence. Anti-Sec17 IgG inhibited Ca^2+^ efflux triggered by preventing trans-SNARE pairing. Fluorescence was measured with an excitation of 485 nm and emission at 520 every 30 seconds for 80 minutes. **(D)** Quantitation of experiments in panel **(C)**. The triangle on the x-axis of panel C represents the time of peak Ca^2+^ release that was used to analyze differences in panel D. Statistical significance was determined by one-way ANOVA for multiple comparisons [F (5, 18) = 100.5; \*\*\*\**p*<0.0001]. Error bars represent mean ± SEM. Tukey’s multiple comparison test was used for individual *p* values (n=3).

### Vacuolar Calcium transport is not altered in *elo3*Δ vacuoles

Another cellular function altered by SL content is Ca^2+^ transport. It has been shown that the lysosphingolipid sphingosine 1-phosphate promotes Ca^2+^ mobilization (Shatrov *et al*, 1997). Others have shown that Ca^2+^ transport by TRP channels can be inhibited by degrading sphingomyelin with sphingomyelinase, extraction of cholesterol with methyl-β-cyclodextrin, or blocking sphingosine biosynthesis with myriocin each of which would increase membrane fluidity (Sághy *et al*, 2015). Furthermore, increased membrane fluidity, as we see with *elo3*Δ vacuoles, has been shown to decrease Ca^2+^ transport by ATPase driven pumps (Grebowski *et al*, 2013; Muzulu *et al*, 1995). Vacuoles take in Ca^2+^ using the high affinity ATPase Pmc1 and the low affinity H^+^/Ca^2+^ antiporter Vcx1 (Cunningham & Fink, 1996, 1994). After Ca^2+^ uptake, vacuoles release the cation upon the formation of *trans*-SNARE complexes between vesicles (Merz & Wickner, 2004). Our previous work used Cal520 fluorescence to show that Ca^2+^ transport by isolated vacuoles was regulated by vacuole acidification and the lipid PI(3,5)P_2_ (Miner *et al*, 2020; Zhang *et al*, 2022b). Here we found that both WT and *elo3*Δ vacuole imported Ca^2+^ upon the addition of ATP (**Fig. 3E, F**). After uptake, both WT and *elo3*Δ vacuoles released Ca^2+^ with the same kinetics and intensity. As a control for SNARE-dependent Ca^2+^ efflux we added antibody against Sec17 to reactions to block SNARE priming and eventual pairing. This blocked Ca^2+^ release by WT and *elo3*Δ vacuoles. Together these indicated that while SL content can influence Ca^2+^ transport in some systems, yeast vacuole Ca^2+^ mobilization does not require C26-VLCFAs.

### The tethering and docking steps are altered on *elo3*Δ vacuoles

In order to narrow when C26-SL were needed for fusion we started by testing SNARE activation by monitoring the release of Sec17 after Sec18-mediated separation of cis-SNAREs into individual coils (Mayer *et al*, 1996). This showed that there was little difference between WT and *elo3*Δ vacuole SNARE priming (not shown). This suggested that the effect of lacking C26-SL was between tethering and hemifusion. To monitor tethering we used fluorescent microscopy as previously described (Eitzen *et al*, 2002; Fratti *et al*, 2004; Wang *et al*, 2002b). Vacuoles isolated from each strain were incubated for 20 min, stained with MDY-64 and mounted on glass slides for visualization. We found that WT vacuoles clustered with a majority of the tethered/docked clusters containing five or more vacuoles (**Fig. 4A-B**). In comparison *elo3*Δ clusters contained fewer vacuoles and approximately 40% of *elo3*Δ were found alone. To specifically inhibit the tethering step, we incubated vacuoles under the same conditions but in the presence of purified Gdi1, a guanine-nucleotide dissociation inhibitor (GDI) that functions in Rab-GTPase turnover by binding to GDP-bound Rab proteins, blocking nucleotide exchange and extracting them from the resident membrane (Garrett *et al*, 1994; Haas *et al*, 1995). We found that adding Gdi1 to WT vacuoles mimicked the docking efficiency of *elo3*Δ vacuoles. Gdi1-resistant clusters were those that have moved onto SNARE pairing. Ungermann et al., showed that tethering required Ypt7 and not SNAREs (Ungermann *et al*, 1998). Therefore, we propose that C26-SL regulated the Ypt7-dependent tethering stage.

**Figure 4.**
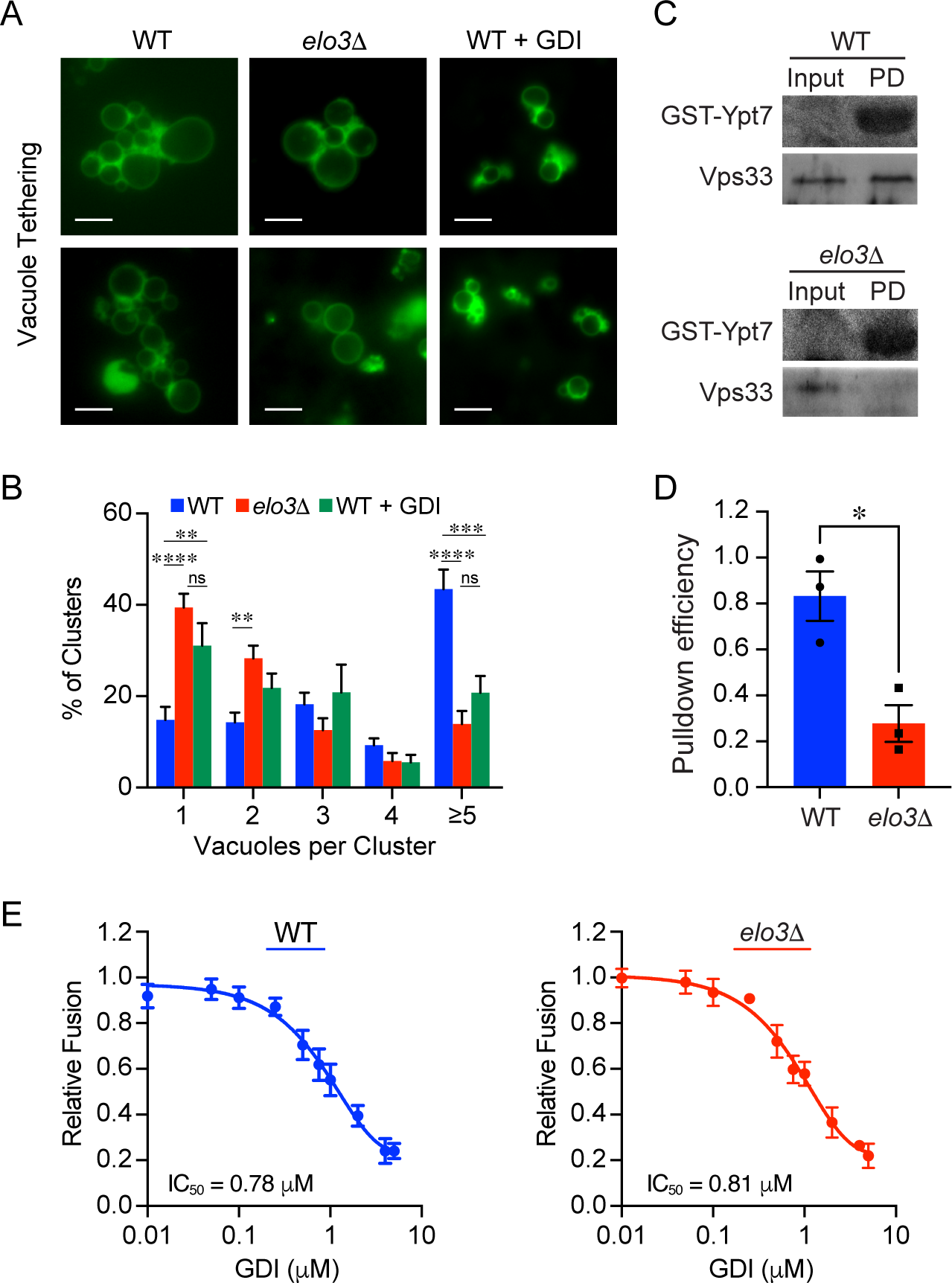
Vacuole tethering is defective in *elo3*Δ vacuoles. **(A)** Isolated vacuoles were incubated for 20 min at 27°C under docking conditions. Following incubation reaction mixtures were placed on ice and gently mixed with the yeast vacuole marker MDY64. Wild-type vacuoles were incubated with purified 2 µM GDI to specifically inhibit the Ypt7-dependent tethering and subsequent docking of vacuoles. **(B)** Quantification of the clustering of docked vacuoles from experiments in (A). Approximately 300 vacuole clusters were counted for each condition in three experiments. Significance was measured using one-way ANOVA for multiple comparisons [F (14, 295) = 11.75, *p*<0.0001]. Error bars represent the mean ± SEM. Tukey’s post hoc multiple comparisons test was used for individual *p* values. Error bars represent mean SEM. (n = 3). ** *p*<0.01; *** *p*<0.001; ns, not significant **(C)** Co-isolation experiments performed with GST-Ypt7 immobilized on glutathione agarose resin and preloaded with GTP. 5X fusion reactions containing vacuoles from WT or *elo3*Δ were solubilized with TX100 and incubated separately with Ypt7-loaded beads. The amount of Vps33 that was bound to resin and found in the input sample were determined by Western blotting. **(D)** Quantification and comparison of the GST-Ypt7 pull-down efficiency. Efficiency was calculated within a strain to correct for the decreased amount of Vps33 on isolated *elo3*Δ vacuoles. (n = 3). Error bars represent Mean ± SEM. Significance was determined using unpaired Student t-test. *P*<0.05. **(E)** Sensitivity of fusion to GDI. Fusion assays using WT and *elo3*Δ vacuoles were performed using a concentration curve of GDI. Fusion was measured at the end of 120 min. Each strain was normalized to its own maximum fusion set to 1. The IC_50_ values were calculated using Graphpad Prism 10. The normalized data was log transformed, normalized and fit to a non-linear regression curve to yield IC_50_ concentrations of GDI for WT and elo3Δ vacuole fusion. Error bars represent the mean ± SEM.

To confirm that the loss of functional Elo3 significantly impaired the tethering state we performed co-affinity isolation assays of GST-Ypt7 and the HOPS complex (Brett *et al*, 2008; Karim *et al*, 2018). To test for Ypt7-HOPS interactions we used recombinant un-prenylated GST-Ypt7 as previously described (Price *et al*, 2000a). GST-Ypt7 was bound to glutathione agarose beads and loaded with GTP. Pull-down experiments were performed with 30 μg of vacuolar material from WT or *elo3*Δ membranes where native GDP-Ypt7 was extracted by GDI. Vacuoles were then solubilized with detergent, mixed with resin bound GST-Ypt7 and incubated for 2h at 4°. After incubation, the unbound material was removed, and resin bound Ypt7 complexes were washed prior to elution with reduced glutathione. Samples were resolved by SDS-PAGE and probed for the presence of Vps33 by Western blotting. We found that Vps33 from WT vacuolar extracts bound to GST-Ypt7 (**Fig. 4C-D**). In contrast the extract from *elo3*Δ vacuoles failed to show an interaction between these proteins. Because the Ypt7 used in these experiments was expressed in bacteria and not prenylated, we can conclude that the altered interactions were not due to changes in Ypt7 conformation or nucleotide binding state. Instead, it appeared that the defect was due to changes in HOPS on *elo3*Δ vacuoles.

Finally, we asked whether the nucleotide binding state of Ypt7 on vacuoles was altered in the absence of C26-SL. To do this we tested for changes in sensitivity to GDI treatment during fusion. As mentioned above, GDI can bind to GDP-bound Ypt7 to promote its extraction from membranes. Thus, changes in GDI sensitivity would suggest that Ypt7 on *elo3*Δ vacuoles were shifted to binding more or less GDP relative to WT, which would differentially affect fusion. To test this, we used a concentration curve of GDI in fusion assays containing WT or *elo3*Δ vacuoles. Both curves were normalized to their own maximum fusion in the absence of GDI. We found that the sensitivity of *elo3*Δ vacuole fusion to GDI was identical to that of WT vacuoles (**Fig. 4E**). This suggests that a portion of Ypt7 on *elo3*Δ vacuoles is functional and that the attenuated fusion in the mutant was due to an overall reduction in tethering events.

### Ypt7 is mislocalized on *elo3*Δ vacuoles

While Ypt7 is equally abundant on WT and *elo3*Δ vacuoles (**Fig. 2H**), the defect in tethering and HOPS interactions suggest that other tethering related events could be dysregulated. During tethering and docking the lipids and proteins that regulate fusion accumulated in a membrane microdomain called the vertex ring (Eitzen *et al*, 2002; Fratti *et al*, 2004; Wang *et al*, 2003b, 2002b). To see if Ypt7 properly localized to the vertex ring, we used vacuoles expressing GFP-Ypt7 under control of its native promoter and performed ratiometric fluorescence microscopy with phosphatidylserine probe PSS380 to label the entirety of the membranes (Wang *et al*, 2002b; Fratti *et al*, 2004). We found that GFP-Ypt7 was enriched at the vertex ring of WT vacuoles as previously reported (**Fig. 5A-B**). When we examined *elo3*Δ vacuoles we found that GFP-Ypt7 failed to accumulate at vertices and contained more at outer membrane edges relative to the WT. This indicated that C26-SL were required for its localization to the vertex. This could lead to fewer interactions with HOPS on *elo3*Δ vacuoles that were needed for optimal tethering.

**Figure 5.**
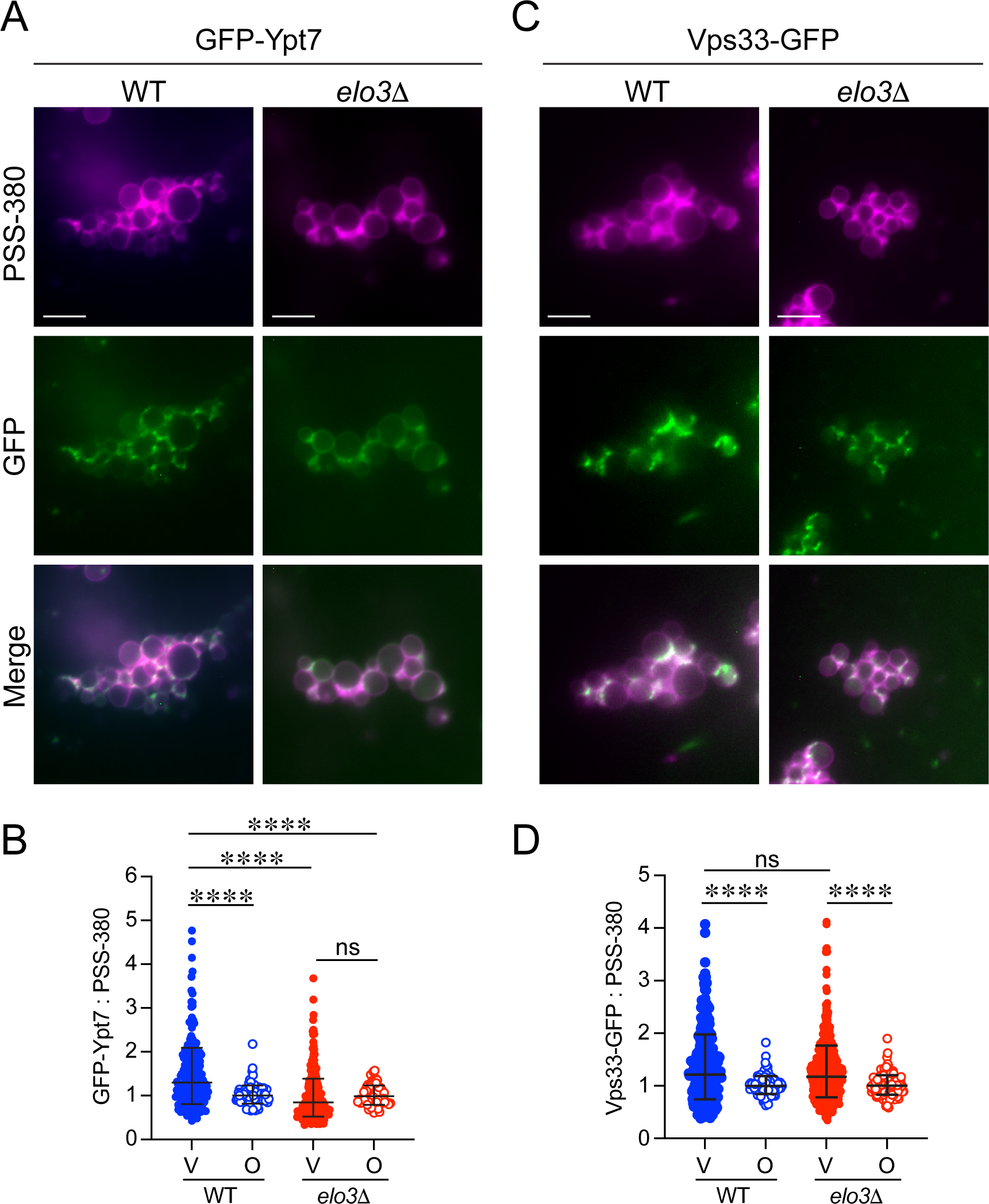
Ypt7 is mislocalized in *elo3*Δ yeast. **(A)** WT and *elo3*Δ vacuoles containing GFP-Ypt7 were incubated under docking conditions for 20 min at 27°C. After incubation, the reactions were placed on ice and labeled with PSS-380. Representative images of docked vacuoles with a line were used for the line scan analyses right of the micrographs. **(B)** Quantitation of ratiometric fluorescence intensities for vertices (V) and outer edge (O) in panel A. The data points were pooled from multiple experiments where each experiment used 15-20 clusters with at least 10 vacuoles per cluster (n>200 vertices for each strain; n>100 outer domain measurements for each strain). Bars represent geometric means ± geometric SD. Significance was measured using one-way ANOVA for multiple comparisons [F (3, 634) = 6.779, *** *p*=0.0002]. Tukey’s post hoc multiple comparisons test was used for individual *p* values. **** *p*<0.0001; ns, not significant. **(C).** WT and *elo3*Δ vacuoles containing Vps33-GFP were analyzed for vertex enrichment as described above. Shown are representative images of docked vacuoles for each strain. **(D)** Quantitation of ratiometric fluorescence intensities for vertices (V) and outer edge (O) in panel C. The data points were pooled from multiple experiments where each experiment used 15-20 clusters with at least 10 vacuoles per cluster (n>400 vertices for each strain; n>200 outer edge measurements for each strain). Bars represent geometric means ± geometric SD. Significance was measured using one-way ANOVA for multiple comparisons [F (3, 1148) = 24.71, *** *p*<0.0001]. Tukey’s post hoc multiple comparisons test was used for individual *p* values. **** *p*<0.0001; ns, not significant.

### The vertex enrichment of HOPS subunits is not altered on *elo3*Δ vacuoles

When we tested the association of HOPS with recombinant GST-Ypt7 we found that HOPS from *elo3*Δ cells failed to interact with the Rab. Coupled with the mislocalization of GFP-Ytp7 on docked vacuoles we predicted that HOPS would also fail to accumulate at vertex sites. To test this, we used WT and *elo3*Δ vacuoles that harbored Vps33-GFP to perform docking assays. We found that unlike the mislocalization of Ypt7, Vps33 enrichment at vertex sites was not affected on *elo3*Δ vacuoles (**Fig. 5C-D**). Vps33 is part of the core Class-C complex that does not directly contact Ypt7. Thus, it was possible that HOPS subunits that do touch Ypt7 could be mislocalized. Both Vps41 and Vps39 directly contact Ypt7 on vacuoles (Wurmser *et al*, 2000; Brett *et al*, 2008; Cabrera *et al*, 2009). Recently, HOPS was found to be assembled from subcomplexes containing Vps11 and Vps39 (HOPS-2) with ones composed of Vps33, Vps16, Vps18 and Vps41 (HOPS-4) (Zhang *et al*, 2023). To see if there was a separation of HOPS subunits on *elo3*Δ vacuoles we monitored GFP-Vps39. Similar to Vps33-GFP, we found that there was no difference in GFP-Vps39 enrichment on WT versus *elo3*Δ vacuoles (not shown). Based on this, we propose that the mislocalization of GFP-Ypt7 was not due to altered HOPS sorting to vertex microdomains. Thus, it is possible that the lack of C26-SL alters the conformation or Ypt7 or HOPS to alter interactions and reduce the vertex enrichment of the Rab.

### PI3P and ergosterol are not affected by the loss of C26-SL

Vertex microdomains are not only enriched in the proteins that drive fusion, but they are also enriched in regulatory lipids including phosphoinositides, DAG and ergosterol (Fratti *et al*, 2004). Importantly, the accumulation of SNAREs, HOPS and Ypt7 depend on these lipids for their collection at vertices. Since C26-SL affect raft-like membrane microdomain structures we examined if regulatory lipids could accumulate at vertex domains of *elo3*Δ vacuoles. To do this we used the tandem FYVE domain from human HRS as a probe for phosphatidylinositol 3-phosphate (PI3P) as previously described (Gillooly *et al*, 2000; Fratti *et al*, 2004). Recombinant GST-FYVE was conjugated to the fluorophore Cy5 and used in vacuole docking reactions at a concentration that does not inhibit fusion. We found that both WT and *elo3*Δ accumulated comparable amounts of Cy5-FYVE at vertex sites (**Fig. 6A-B**), suggesting that C26-SL were not required for the accumulation of lipids at vertex microdomains.

**Figure 6.**
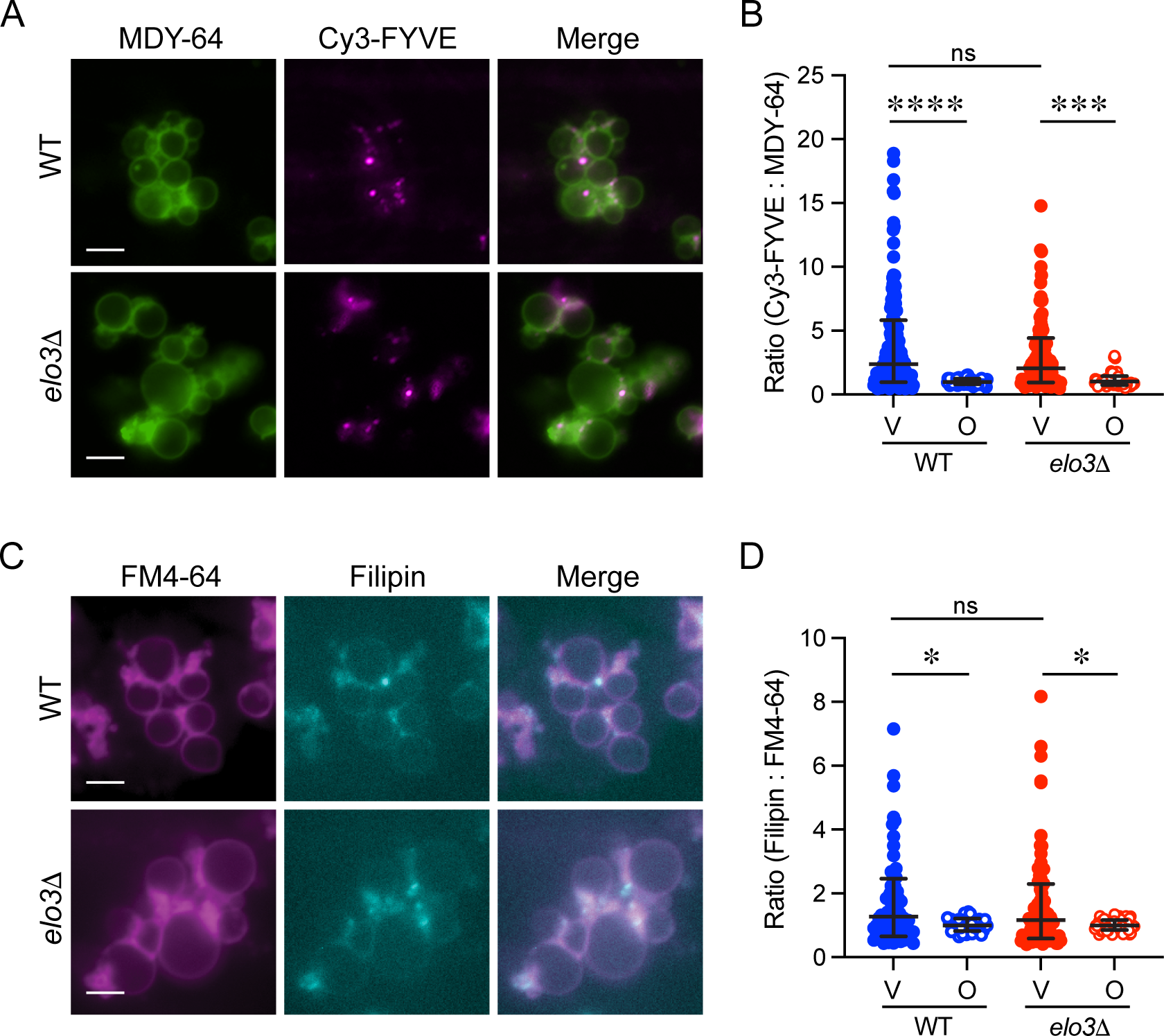
Vertex enrichment of PI3P and ergosterol does not require C26-SL. **(A)** Docking reactions containing 6 μg of purified vacuoles were performed in the presence of 0.25 μM Cy5-FYVE. Yeast Vacuole Marker MDY-64 was added at the end of the reaction and docked vacuoles were visualized using fluorescence microscopy. **(B)** Quantitation of ratiometric fluorescence intensities for vertices (V) and outer edge (O) in panel A. The data points were pooled from multiple experiments where each experiment used 15-20 clusters with at least 10 vacuoles per cluster (n>400 vertices for each strain; n>200 outer edge measurements for each strain). Bars represent geometric means ± geometric SD. Significance was measured using one-way ANOVA for multiple comparisons [F (3, 452) = 24.71, *** *p*<0.0001]. Tukey’s post hoc multiple comparisons test was used for individual *p* values. **** *p*<0.0001; *** *p*<0.001 ns, not significant. **(C)** Docking reactions performed in the presence of 5 μM filipin III. FM4-64 was added at the end of the reaction and docked vacuoles were visualized by fluorescence microscopy. **(D)** Quantitation of ratiometric fluorescence intensities for vertices (V) and outer edge (O) in panel C. The data points were pooled from multiple experiments where each experiment used 15-20 clusters with at least 10 vacuoles per cluster (n>100 vertices for each strain; n>50 outer edge measurements for each strain). Bars represent geometric means ± geometric SD. Significance was measured using one-way ANOVA for multiple comparisons [F (3, 291) = 6.152, *** *p*=0.0005]. Tukey’s post hoc multiple comparisons test was used for individual *p* values. * *p*<0.05; ns, not significant.

The formation of membrane lipid rafts or nanodomains in yeast membranes is strongly influenced by the amount of sphingolipids and ergosterol (Klose *et al*, 2010; Toulmay & Prinz, 2013). Other studies have shown that ergosterol concentrates at vertex regions during vacuolar fusion and sterol-enriched vacuoles show enhanced fusion (Fratti *et al*, 2004; Tedrick *et al*, 2004). We next asked if the lack of C26-SL would affect the vertex localization of ergosterol. Using subinhibitory amounts of the fluorescent macrolide filipin III, which binds to ergosterol, we compared the fluorescence of WT and *elo3*Δ docked vacuoles (**Fig. 6C-D**). When the docking reactions of WT vacuoles were visualized, we observed strong fluorescence as puncta at the vertex and at the boundary domains. When vacuoles isolated from *elo3*Δ cells were incubated with filipin III we found no significance difference in vertex enrichment between WT and *elo3*Δ. Altogether, the data suggests that yeast sphingolipids containing C26-SL do not strongly affect the localization of other regulatory lipids. While ergosterol and PI3P localization are not affected on elo3Δ vacuoles, it is still possible that a physical aspect of these membranes, e.g. fluidity, is sufficiently altered to affect Ypt7 localization, which ultimately reduces the fusion efficiency of these vacuoles.

Finally, we asked if GFP-Ypt7 localization would be affected *in vivo* when sphingolipid production was inhibited. To do this we treated WT cells expressing GFP-Ypt7 with Aureobasidin A (AbA) and compared Ypt7 distribution in *elo3*Δ cells. In logarithmically growing cells GFP-Ypt7 was evenly distributed around the vacuole to form complete rings that overlap with FM4-64 (Wang *et al*, 2002b; Sasser *et al*, 2012a). We observed the same rings here, however, when cells were treated with AbA we found that GFP-Ypt7 formed fewer rings and with reduced intensity suggesting that SL production was needed for normal Ypt7 distribution (**Fig. 7A-B**). The altered distribution of Ypt7 in WT treated with AbA was indistinguishable from the distribution of Ypt7 in *elo3*Δ further linking SL production and Ypt7. In addition to rings, Ypt7 in whole cells also collects in punctate domains that overlap with FM4-64 with some found within the vacuole lumen. The latter was likely due to post-fusion internalization of membrane fragments (Karim *et al*, 2018). Unlike the differences seen in ring formation between WT, *elo3*Δ and AbA-treated cells, we did not observe differences in puncta formation (**Fig. 7A and C**). Based on the role of SL in lipid raft formation and what we observed when we measured membrane fluidity, we propose that SL normally provide membrane stiffness needed for Ypt7 to localize where it functions during membrane fusion. In other words, vacuoles do not function properly when membranes are too fluid.

**Figure 7.**
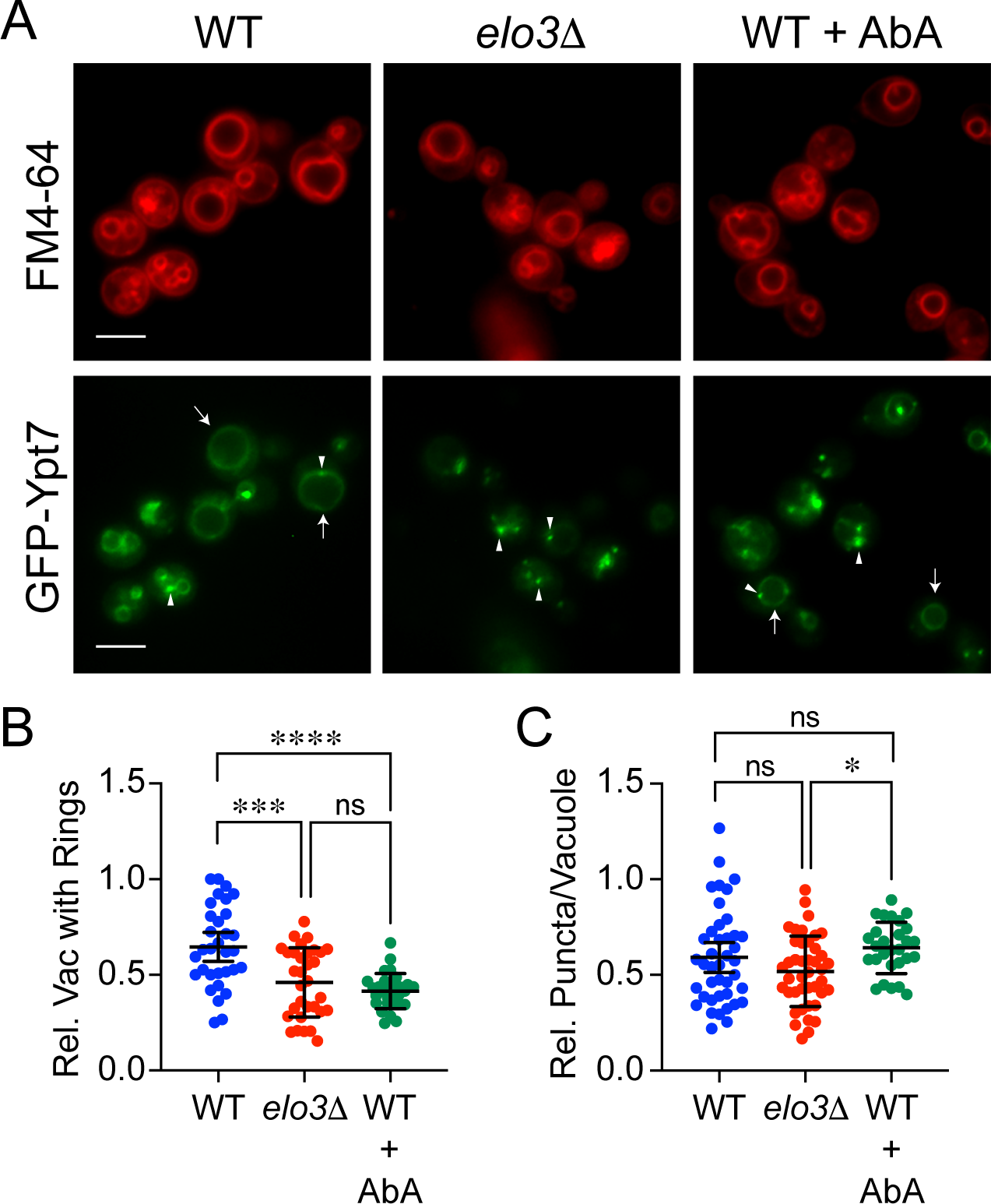
In vivo GFP-Ypt7 localization in altered in cells with inhibited SL production. **(A)** Wild type and *elo3*Δ yeast expressing GFP-Ypt7 were grown overnight in selective media and back diluted to ∼0.7 OD_600_ the next day. Cells were transferred to YPD broth and incubated for 4 h at 30°C while shaking. WT cells were also treated with AbA. Cells were then incubated with 10 µM MDY-64 for 15 min in the dark before visualization. Images are representative of three separate trials. Arrows represent complete rings of GFP-Ypt7 around a vacuole. Arrow heads represent GFP-Ypt7 puncta. Scale bars: 4 μm. **(B)** Quantitation of the number of vacuoles with complete rings of GFP-Ypt7 per cell. Ten 63x fields were counted for a total of 200-600 vacuoles per condition/strain for each experiment. The number of vacuoles with complete GFP-Ypt7 in each field was converted to a percentage of total amount of vacuoles. Bars represent mean ± 95% CI. Significance was measured using one-way ANOVA for multiple comparisons [F (2, 90) = 16.00, **** *p*<0.0001]. Tukey’s post hoc multiple comparisons test was used for individual *p* values. *** *p*<0.0001, **** *p*<0.0001; ns, not significant **(C)** Quantitation of complete GFP-Ypt7 puncta per vacuole. Ten 63x fields were counted for a total of 100-300 puncta per vacuole for each experiment. The number of puncta per vacuole in each field was converted to a percentage of total amount of vacuoles. Significance was measured using one-way ANOVA for multiple comparisons [F (2, 108) = 3.321, *p*=0.0398]. Bartlett’s post hoc multiple comparisons test was used for individual *p* values. * *p*<0.05; ns, not significant

## Discussions

Studies have shown that C26 VLCFAs are found almost exclusively within sphingolipids at the amide bond position, leaving only trace amounts of free VLCFAs within the cell (Lester *et al*, 1993; Megyeri *et al*, 2016). Therefore, we posit that the observations outlined above stem from a lack of C26-SL, as opposed to a lack of free cerotic acid. This is bolstered by the observation that cells treated with sphingolipid biosynthesis inhibitors display aberrant vacuolar morphology similar to *elo3*Δ cells. WT yeast produce sphingolipids that most often (70-90%) contain a C26:0:1 (cerotic acid with no double bonds, and a single hydroxylation within the acyl chain), whereas yeast that lack Elo3 have been shown to produce non-canonical sphingolipids with shorter acyl chains (20-24 carbons). This is accompanied by a significant decrease in complex sphingolipid levels (Ejsing *et al*, 2009; Oh *et al*, 1997). In contrast, the bulk of the glycerol-based phospholipids, whose fatty acid chains are usually 16 or 18 carbons long, and typically contain one or two desaturated fatty acids (16:1 and/or 18:1) such as PI3P were not affected by the absence of C26-SL on the vacuole. Importantly, we found that yeast strains lacking C26 VLCFAs are unable to enrich the Rab Ypt7 at the vertex region and fail to support its interactions with HOPS, which is likely due to increased membrane fluidity.

The observation that vacuoles isolated from cells lacking C26-SL are fusion deficient could arguably be attributed to either one or both of the following: biophysical requirements of the membrane – such as interdigitation of acyl chains, or strong hydrogen bonding (Haque *et al*, 2001; Peter Slotte, 2013); and/or sphingolipid-rich nanodomains at vertex regions that serve as platforms for fusion protein concentration and localized activity (Harayama & Riezman, 2018; Raghupathy *et al*, 2015; Sezgin *et al*, 2017). Others have found that VLCFAs can stimulate membrane fusion in plant cells during cytokinesis and with liposomes (Molino *et al*, 2014). This is consistent with the inhibition of fusion when VLCFAs are absent.

It was initially shown that lipids isolated from yeast can form into model membranes that separate into visibly distinct domains when they were labeled with fluorescent lipophilic dyes with different lipid-partitioning properties (Klose *et al*, 2010). The domain that was found to be liquid-ordered-like (L_o_-like) was enriched in BODIPY-cholesterol, and this domain dissipated and was visually absent when the lipids were isolated from yeast lacking a functional Elo3. This phase separation in yeast lipids was demonstrated again when it was shown that when yeast cells are grown to stationary phase the vacuole membrane displays regions that separate into distinct lipid domains *in vivo* (Toulmay & Prinz, 2013; Reinhard *et al*, 2023; Leveille *et al*, 2022). These domains were found to be enriched in either Vph1 or Ivy1 (a PI(3,5)P_2_-binding protein that interacts with Ypt7 and Vps33 (Lazar *et al*, 2002; Malia *et al*, 2018; Numrich *et al*, 2015), and they display properties that are indicative of differences in relative membrane fluidity. Fluorescently tagged Vph1 colocalizes with a dye that partitions into liquid-disordered-like (L_d_-like) membranes, while a fluorescent Ivy1 construct colocalizes with the macrolide filipin III, which binds to ergosterol in yeast membranes in the L_o_-like region of the membrane. Vph1 localization was subsequently utilized as a negative marker in studies to show that the regulatory fusion lipids PI3P, PI4P, and PI(3,5)P_2_ tend to enrich in regions that lack Vph1 and intramembrane particles (IMPs) (Takatori *et al*, 2016; Tomioku *et al*, 2018), *i.e.* these known regulatory lipids enrich in the raft-like domains that are also known to contain relatively high amounts of sphingolipids and sterols. However, it should be noted that other studies have demonstrated an activation of the V-ATPase upon the addition of exogenous, short-chain PI(3,5)P_2_ and a stabilization of the V_O_-V_1_ complex by this lipid, suggestive of a protein-lipid interaction (Li *et al*, 2014) and the possibility of separate pools of PIP lipids.

One possibility for the lack of this interaction could be linked to HOPS phosphorylation by the casein kinase Yck3. This would be consistent with other studies showing that casein kinases are activated by sphingolipids (Gavrilova *et al*, 1995; Nielander *et al*, 1995). In rat synaptosomes, casein kinase 2 associates with detergent resistant microdomains and its activity is abolished when cholesterol is extracted (Gil *et al*, 2011). Thus, it is possible that Yck3 is not fully active on *elo3*Δ vacuoles leading to a disruption in the binding of HOPS and Ypt7. Zick and Wickner found that HOPS phosphorylation (P-HOPS) by Yck3 alters binding to Ypt7 (Zick & Wickner, 2012). Specifically, they found that P-HOPS preferentially binds GTP-loaded Ypt7 versus GDP-Ypt7; however, the overall binding of P-HOPS to Ypt7 was sharply reduced when compared to Ypt7 binding by unphosphorylated HOPS. This suggests that Yck3 in general inhibits HOPS binding to Ypt7. This is in keeping with reports by LaGrassa *et al*. showing that phospho-Vps41 does not bind GDP-Ypt7 (LaGrassa & Ungermann, 2005). Other data further suggests that Yck3 phosphorylation of HOPS is not responsible for the mislocalization of Ypt7 on *elo3*Δ vacuoles. When *YCK3* is deleted, vacuole fusion becomes resistant to GDI and the GTPase activating factor Gyp7 (LaGrassa & Ungermann, 2005; Brett *et al*, 2008). We found that the IC50 of GDI was the same for WT and *elo3*Δ vacuoles, indicating that Yck3 function does not play a role here.

## Materials and Methods

### Yeast strains and growth conditions

Yeast strains used in this study are found in Table 1. Yeast strains were grown in YPD (1% yeast extract [RPI Y20020], 2% peptone [RPI P20240], 2% dextrose [RPI G32040], or synthetic dropout media (YNB, yeast nitrogen base lacking amino acids and ammonium sulfate [RPI Y20060], with amino acids, adenine sulfate [RPI A12575], and uracil (Millipore-Sigma U0750) added at ∼76 mg/L, except leucine which was at ∼380 mg/L (RPI L22000) lacking the specified nutrient the specified nutrient. The *ELO3* coding sequence was deleted using homologous recombination by amplifying the *hphMX6* gene conferring hygromycin resistance from the vector pAG32 (Addgene 35122) using the primers elo3F (5’-CGGCTTTTTTCCGTTTGTTTACGAAACATAAACAG-TCATCTGTTTAGCTTGCCTTGTCC-3’) and elo3R (5’-TTTTTTCTTTTTCATTCGCTGTCAAAAATTCTCGCTTCCGACACTGGATGGCGGCGTTA-3’) (IDT). The PCR product designed for homologous recombination was transformed using lithium acetate (Sigma L6883) as previously described (Tripp *et al*, 2013), and cells were grown on YPD agar plates containing 200 mg/L hygromycin B (Goldbio H-270-2-1). All enzymes used for DNA manipulation and subcloning were purchased from New England Biolabs. The construct was confirmed via sequencing (ACGT). All plasmids were transformed into wild type and *elo3*Δ strains via electroporation (BIORAD MicroPulser Electroporator 1652100) and plated onto appropriate auxotrophic selection plates.

**Table 1.** Strains used in this study.

| **Strain** | **Genotype** | **Source** |
| --- | --- | --- |
| BJ3505 | *MATa pep4::HIS3 prb1-Δ1.6R his3-200 lys2-801 trp1Δ101 (gal3) ura3-52 gal2 can1* | (Jones *et al*, 1982) |
| DKY6281 | *MATα leu2-3,112 ura3-52 his3-Δ200 trp1-Δ901 lys2-801 suc2-Δ9 pho8Δ::TRP1* | (Klionsky & Emr, 1989) |
| SEY6210 | *MATα leu2-3,112 ura3-52 his3-Δ200 trp1-Δ901 lys2-801 suc2-Δ9* | (Klionsky & Emr, 1989) |
| GFP-Ypt7 | *SEY6210, ypt7::HIS3 pRS304:GFP-Ypt7 (TRP1)* | (Wang *et al*, 2002b) |
| RFY120 | *DKY6281, elo3::hphMX6* | This study |
| RFY121 | *BJ3505, elo3::hphMX6* | This study |
| RFY124 | *SEY6210, ypt7::URA3 elo3::hphMX6 pRS304:GFP-Ypt7* | This study |
| Vps33-GFP | *DKY6281, VPS33-GFP* | (Wang *et al*, 2002b) |
| RFY125 | *DKY6281, elo3::hphMX6 VPS33-GFP* | This study |
| GFP-Vps39 | *SEY6210, vps39::HIS3 pYIPlac211-GFP-VPS33 (TRP)* | (Wang *et al*, 2002b) |
| RFY126 | *SEY6210, vps39::HIS3 elo3::hphMX6 pYIPlac211-GFP-VPS33* | This study |

### Live microscopy

General vacuolar morphology of live yeast was determined by growing cells overnight in YPD at 30°C to mid-log phase. Cells were then diluted in fresh YPD to OD_600_ ∼ 0.6 and allowed to grow for 1 h, before treatment with 62.3 μM myriocin (Cayman 63150), 113 nM aureobasidin A (Takara 630466), 50 μM cerulenin (Cayman 10005647), or methanol (Macron) vehicle control. After additional growth in the presence of inhibitor for ∼5 h, Yeast Vacuole Membrane Marker MDY-64 (Thermo Fisher Y7536) was added to a final concentration of 10 µM, and cells were incubated at 30°C for 15 min. Cells were then isolated by centrifugation (12000 x g, 1 min, RT) and resuspended in PBS. Twenty microliters of this cell suspension were then mixed 1:1 with 0.6% low-melt agarose (RPI A20070) warmed to 50°C, and 10 µL of the mixture was placed on a glass slide (AmScope BS-144P-200S-22) and covered with a glass coverslip (Thermo) for viewing by fluorescence microscopy. For strains expressing a fluorescent construct, overnight growth was in SC dropout media for plasmid maintenance before cells were diluted into YPD the day of visualization. For strains expressing a green fusion protein, the cells were grown in the presence of 10 µM FM4-64 (Thermo Fisher T-3320) for the ∼6 h following back dilution, while those that expressed red fusion proteins were incubated with 10 µM MDY-64 for the final 20 min prior to visualization. Images were acquired using a Zeiss Axio Observer Z1 microscope equipped with an X-Cite 120XL light source, Plan Apochromat x63 oil objective (NA 1.4) and an AxioCam CCD camera. All images were analyzed with ImageJ software (NIH). FM4-64 images were visualized with a 42 HE CY 3 shift-free filter set, Filipin was visualized with a 49 DAPI shift free filter set and GFP was visualized using a 38 HE EGFP shift-free filter set.

### Vacuole isolation and in vitro content mixing

Yeast vacuoles were isolated as described (Haas *et al*, 1994) with slight modifications. Cells were grown at 30°C and shaking at 225 rpm in 1L YPD broth in a 2L Erlenmeyer flask to OD_600_ values of 0.7-1.1. Cells were then pelleted by centrifugation (3000 x g, 5 min, 24°C) and washed with 50 mL of 100 mM TRIS-Cl (TRIS base, RPI T60040), pH 9.4, 15 mM DTT (RPI D11000), and incubated at 30°C for 10 min. The cells were pelleted again as above and resuspended in 15 mL spheroplasting buffer (50 mM potassium phosphate buffer pH 7.5, and 600 mM sorbitol in YPD containing 0.2% dextrose) with 0.8-1.5 mg/mL oxalyticase per O.D. unit of cells. Recombinant oxalyticase was purified as described (Scott & Schekman, 1980). Cells were incubated for 30-45 min at 30°C and transferred to chilled oak ridge tubes before spinning for 10 min at 1000 x g 4°C. All other steps in vacuole isolation were identical as referenced. Content mixing reactions (30 μL) contained 3 μg each of vacuoles from BJ3505 (*pep4*Δ *PHO8)* and DKY6281 (*PEP4 pho8*Δ) backgrounds in fusion reaction buffer consisting of PIPES-sorbitol (PS) buffer (20 mM PIPES-KOH [RPI P40140], pH 6.8, 200 mM sorbitol [RPI S23080]), 125 mM KCl (RPI P41000), 5 mM MgCl_2_ (RPI M25000), ATP regenerating system (1 mM ATP [RPI A30030], 0.1 mg/mL creatine kinase [Sigma 10127566001], 29 mM creatine phosphate [Gold Biotech C-323-5]), 10 μM CoA (Sigma C4282), and 280 nM recombinant Pbi2 (IB2, Protease B inhibitor)(Slusarewicz *et al*, 1997). Reactions were incubated at 27°C and Pho8 activity was assayed in 250 mM Tris-Cl, pH 8.5, 0.4% Triton X-100 (RPI 400001), 10 mM MgCl_2_, 1 mM *p*-nitrophenyl phosphate (MP Biomedicals 0210087810). Fusion was monitored as the absorbance at 400 nm from *p-*nitrophenolate production through phosphatase activity.

### Lipid mixing

Lipid mixing assays to track outer leaflet fusion (hemifusion) were performed as previously (Qiu & Fratti, 2010; D’Agostino & Mayer, 2019). Vacuoles were isolated from BJ3505 WT and BJ3505 *elo3*Δ yeast, and 300 μg of vacuoles (by protein) were next labeled by incubating with 400 µL of a 150 μM rhodamine B DHPE (1,2-dihexadecanoyl-sn-glycero-3-phosphoethanolamine) (RhPE/Lissamine rhodamine; ThermoFisher L1392) in PS buffer while nutating for 10 min at 4°C. Following labeling, 800 μL of 15% Ficoll PM 400 (Cytiva 17030050) (diluted in PS buffer) was mixed with the vacuole-RhPE mixture and transferred to 11 x 60 mm polyallomer ultracentrifuge tube, overlaid with 1.2 mL of 8% Ficoll, 1.2 mL of 4% Ficoll, and 0.5 mL of PS buffer (0% Ficoll). Vacuoles were separated from free dye by centrifugation (100,000 x g, 25 min, 4°C, SW-60 Ti rotor, Beckman L8-80M Ultracentrifuge) and recovered from the 0%-4% interface. Lipid mixing reactions (90 μL) contained 2 μg of labeled vacuoles and 16 μg of unlabeled vacuoles in reaction buffer. Reaction mixtures were transferred to a black, half-volume, 96-well flat-bottom microtiter plate (Greiner Bio-One 675076), and fluorescence from dequenching was monitored (λ_ex_=544 nm, λ_em_=590 nm) in a POLARstar Omega BMG Labtech plate reading fluorometer. Readings were taken every minute for 60 min at 27°C. At the end of incubation each well was treated with 1% TX-100 to solubilize membranes to reach the maximum fluorescence per well. Dequenching due to fusion was calculated as a percentage of the maximum fluorescence at each time point. The data was then normalized to the starting fluorescence set to 1.

### Vacuole acidification

V-ATPase acidification of vacuoles was performed as previously described (Müller *et al*, 2003; Zhang *et al*, 2022a, 2022b). *In vitro* acidification reactions (60 µl) contained 20 µg BJ3505 background vacuoles, reaction buffer, ATP regenerating system, 10 µM CoA, 283 nM Pbi2, and 15 µM acridine orange (Sigma 65-61-2). Reactions were pipetted into a nonbinding black half-volume 96-well flat-bottom plate. ATP regenerating system or PS buffer was added, and reactions were incubated at 27°C. Acridine orange fluorescence was measured in a fluorescence plate reader (POLARstar Omega, BMG Labtech) with the excitation at 485 nm and emission at 520 nm. Reactions were initiated with the addition of ATP regenerating system following the initial measurement. After decreases in fluorescence plateaued (400-600 sec), we added 30 µM FCCP (Carbonyl cyanide-4-(trifluoromethoxy) phenylhydrazone) (Cayman Biochemical 15218) to collapse the proton gradient and restore acridine orange fluorescence. Each trace was normalized to the fluorescence intensities at the time of adding ATP and set to 1.

### Calcium transport

Vacuolar Ca^2+^ transport was measured as described (Miner *et al*, 2020; Miner & Fratti, 2019; Sasser *et al*, 2012b; Zhang *et al*, 2022a). *In vitro* Ca^2+^ transport reactions (60 µl) contained 20 µg vacuoles from BJ3505 backgrounds, reaction buffer, 10 µM CoA, 283 nM Pbi2, and 150 nM of the Ca^2+^ probe Cal520 dextran conjugate MW 10,000 (AAT Bioquest 20601). Reaction mixtures were loaded into a black, half-volume 96-well flat-bottom plate with nonbinding surface. ATP regenerating system was added, and reactions were incubated at 27°C. Reactions were analyzed using a fluorescence plate reader with the excitation filter at 485 nm and emission filter at 520 nm. Samples lacking ATP were used a negative control for Ca^2+^ uptake. Antibody against the SNARE co-chaperone Sec17 was added as a negative control for SNARE dependent Ca^2+^ efflux (Merz & Wickner, 2004). Anti-Sec17 was prepared as described (Mayer *et al*, 1996). Fluorescence readings were taken every 20 seconds for 60 min and normalized to the initial fluorescence value set to 1 for each reaction. Calibration was done using buffered Ca^2+^ standards (Invitrogen, Waltham, MA).

### Tethering and Docking

Isolated vacuoles were subjected to docking assays as previously described (Fratti *et al*, 2004; Wang *et al*, 2002b) with slight modifications. Tethering reactions (30 μL) contained 6 μg of vacuoles isolated from the indicated strains in fusion reaction buffer modified for docking conditions (PS buffer, 100 mM KCl, 0.5 mM MgCl_2_, 0.33 mM ATP, 13 mM creatine phosphate, 33 μg/mL creatine kinase, 10 µM coenzyme A, and 280 nM IB_2_). Tethering was monitored by counting the number of vesicles per cluster using WT or *elo3*Δ vacuoles. As a control for Ypt7 dependent tethering we used recombinant GDI (GDP dissociation inhibitor) to extract GDP-bound Ypt7 to block further tethering. Recombinant GDI with a C-terminal chitin binding peptide was produced as described and dialyzed against 125 mM KCl in PS buffer. (Starai *et al*, 2007). A percentage of isolated vacuoles become tethered during the preparation so complete cluster disruption is not possible.

### Vertex microdomain formation

Vertex enrichment of factors during tethering and docking were performed with vacuoles from cells expressing GFP fusion proteins or labeled with lipid binding probes. To track GFP-Ypt7 localization reactions were incubated under docking conditions as described above and stained with 4 μM PSS-380 or FM4-64 prior to examination (Wang *et al*, 2003b; Fratti *et al*, 2004). PSS-380 labels the bulk lipid phosphatidylserine (a gift from Drs. Smith and Koulov, Notre Dame University, South Bend IN) (Koulov *et al*, 2003). Ergosterol vertex enrichment was monitored with the fluorescent polyene macrolide Filipin III (5 μM) (Cayman 70440) and stained with FM4-64. To localize the distribution of PI3P on vacuoles, reactions were incubated with the PI3P binding protein domain FYVE conjugated to the fluor Cy5 (0.3 µM) and the non-specific membrane dye MDY-64 (0.5 µM). Recombinant GST-FYVE was produced as described and dialyzed against 125 mM KCl in PS buffer (Gillooly *et al*, 2000) and conjugated to mono-reactive NHS ester Cy5 dye following manufacturer’s instructions (Cytiva PA15101). Docking reactions were incubated with Filipin or Cy3-FYVE as previously described (Fratti *et al*, 2004; Sasser *et al*, 2012a). Following incubation at 27°C for 20 min, reactions were mixed with 30 μL of 0.6% low melt agarose in PS buffer melted at 50°C, and 15 μL aliquots were mounted on pre-chilled slides and observed by fluorescence microscopy. Images were acquired using a Zeiss Axio Observer Z1 microscope equipped with an X-Cite 120XL light source, Plan Apochromat x63 oil objective (NA 1.4) and an AxioCam CCD camera. Images were acquired at 23°C without pixel binning. GFP images were acquired using 38 HE EGFP shift-free filter set. Cy3 and FM4-64 images were taken using a 43 HE CY3 shift-free filter set. Cy5 images were taken using a 50 HE CY5 shift-free filter set. PSS380 and filipin images were acquired using a 49 DAPI shift-free filter set. Exposure times were set using WT vacuoles for each fluorescence channel and scripts acquired non-specific images followed by specific reporters. This ensures that bleaching is consistent to negate it as a factor in calculating intensity ratios. Exposure times were held constant within an experiment.

Images were analyzed using ImageJ software (NIH). Vertex enrichment was determined by first measuring maximum fluorescence intensity in each channel at each contact point between membranes, i.e. a vertex domain within a cluster. Next, fluorescence intensity was measured in each channel at outer membrane domains where vacuoles are not in contact with other membranes. The ratio of specific (*e.g.,* GFP) to non-specific (*e.g.,* FM4-64) was determined for vertices and outer membrane domains and compared for relative enrichment. Measurements for each condition were taken of 15-20 clusters to yield 100-300 vertices for each condition/strain per experiment. Data from multiple experiments are combined in column plots showing individual values as well as the geometric means and geometric standard deviation for each condition. One-way ANOVAs and Tukey’s post-hoc analysis for multiple comparisons were used to measure significance. P-values <0.05 were considered significant.

### GST-Ypt7-HOPS Interactions

GST-Ypt7 was purified and loaded onto Pierce glutathione agarose (Thermo) for co-immunoprecipitation analyses as previously described (Brett *et al*, 2008; Price *et al*, 2000a) with minor modifications. *E. coli* BL21(DE3) cells (New England Biolabs C2527I) were grown in auto-induction media (11.33 g Na_2_HPO_4_-7H_2_O dibasic, 3 g KH_2_PO_4_, 20 g tryptone, 5 g yeast extract, 5 g NaCl in per liter) for 18 h, at 37°C, with shaking at 220 rpm (Studier, 2014). Cells were harvested by centrifugation, washed in cold wash buffer (50 mM Tris-Cl, pH 8, 150 mM NaCl, 0.01% 2-mercaptoethanol, 5 mM MgCl_2_, 1 mM PMSF [RPI P20270], and 5 μM leupeptin hemisulfate [Cayman Chemical 14026]), lysed by sonication, and the lysate was separated from debris by ultracentrifugation (Type 70 Ti fixed angle rotor, 200,000 x g, 30 min, 4°C). The cleared lysate was loaded onto glutathione agarose resin (ThermoFisher 16101), washed with wash buffer 10 column volumes of wash buffer, and GST-Ypt7p was eluted with one resin-bed volume of elution buffer composed of wash buffer containing 10 mM reduced glutathione (Sigma G6013).

Following purification, 15 μg of GST-Ypt7p was loaded onto glutathione resin and charged with GTP (Sigma G8877) as previously described (Price *et al*, 2000a). Separately, 6X fusion reactions containing vacuoles isolated from WT or *elo3*Δ yeast were incubated with 2 μM recombinant GDI for 5 min at 27°C to extract resident GDP-bound Ypt7 from vacuoles, reactions were then spun down and resuspended in 180 μL solubilization buffer (25 mM HEPES-KOH, pH 7.5, 150 mM NaCl, 10% glycerol, 0.5% Triton X-100, 5 mM MgCl_2_, 0.01% 2-mercaptoethanol, 1 mM PMSF, 5 μM leupeptin), and 10% was taken as an input from each sample and immediately boiled in SDS-PAGE loading buffer. The remainder was mixed with GTP-Ypt7-loaded glutathione-resin and nutated for 2h at 4°C. Beads were washed 3X with solubilization buffer and eluted by boiling in SDS-PAGE loading buffer. Samples were submitted to SDS-PAGE and immunoblotting for GST-Ypt7 and the HOPS subunit Vps33.

### Membrane fluidity assay

Merocyanine 520 (MC520) was purchased from Sigma (323756) and used as previously described (Barker & Kennedy, 2017; Langner & Hui, 1999; Stillwell *et al*, 1993). Briefly, vacuoles isolated from BJ3505 WT or *elo3*Δ cells (25 µg total protein diluted to 50 µL in PS buffer, 0.5 mg/mL final) were mixed with 6 µL of 100 µM MC520 in PS buffer (Stock MC540 was at 125 mM in DMSO) for a total volume of 100 µL and a final concentration of 0.2% DMSO. These mixtures were incubated in the dark for 20 min before re-isolating the vacuoles by centrifugation (5000 x g for 5 min at room temp) and resuspended in 100 µL of PS buffer containing 2 mM ATP, 2 mM MgCl2, 125 mM KCl and 10 µM CoA. Resuspended mixtures were put in a black half-volume, flat-bottom 96-well plate and endpoint fluorescent emission was measured (λex= 520 nm, λem = 580 nm, CLARIOStar BMG Labtech). Samples of isolated vacuoles were blanked against a reaction containing 6 µL 100 µM MC540 in PS buffer and 50 µL PS buffer. Separate samples were treated with 1% Triton X-100 to solubilize membranes to release MC540 to determine maximum fluorescence at 594 nm.

### Statistical analysis

Results were expressed as the mean ± SEM, mean ± 95% confidence interval (CI), or geometric mean ± SD as needed. Experimental replicates (n) is defined as the number of separate experiments. Statistical analysis was performed by Unpaired two-tailed t-test or One-Way ANOVA for multiple comparisons using Prism 9 (GraphPad, San Diego, CA). Statistical significance is represented as follows: \**p*<0.05, ** *p*<0.01, *** *p*<0.001, **** *p*<0.0001. Tukey or Dunnett post hoc analysis was used for multiple comparisons and individual p-values.

## Data availability

All data generated or analyzed during this study are included in this published article.

## Acknowledgements

The authors wish to thank Dr. Lois Weisman, Dr. Gary Eitzen, and Dr. William Wickner for the generous gifts of antibodies. We also Dr. Alexey Merz for the gift of the pGST-Ypt7-Parallel-1 plasmid. This research was supported by grants from the Natural Science and Engineering Council of Canada (RGPIN/2017-06652) to CLB, and National Institutes of Health (R01-GM101132) and National Science Foundation (MCB 1818310, MCB 2216742) to RAF. J.D.C. was partially supported by an NIGMS-NIH Chemistry-Biology Interface Training Grant (5T32-GM070421).

## Author contributions

**Chi Zhang**: Conceptualization; data curation; formal analysis; investigation; visualization; methodology; writing – original draft; writing – review and editing.

**Logan R. Hurst**: Conceptualization; data curation; formal analysis; investigation; visualization; methodology; writing – original draft; writing – review and editing.

**Zeynep D. Gokbayrak**: Data curation; formal analysis; visualization; writing – review and editing.

**Jorge D. Calderin**: Data curation; visualization; writing – review and editing.

**David A. Rivera-Kohr**: Data curation; visualization; writing – review and editing.

**Adam Balutowski**: Data curation; visualization; writing – review and editing.

**Michael R. Hrabak**: Data curation; visualization; writing – review and editing.

**Thomas D.D. Kazmirchuk**: Data curation; visualization; writing – review and editing.

**Christopher L. Brett**: Conceptualization; formal analysis; supervision; funding acquisition; investigation; visualization; methodology; writing – original draft; writing – review and editing.

**Rutilio A Fratti**: Conceptualization; formal analysis; supervision; funding acquisition; investigation; visualization; methodology; project administration; writing – original draft; writing – review and editing.

## Abbreviations

AbA: Aureobasidin A
C26-SL: 26 carbon VLCFA containing SL
DAG: diacylglycerol
FAS: fatty acid synthase
HOPS: homotypic fusion and protein sorting
IC50: half-maximal inhibitor concentration
IPC: inositol-phosphoceramide
LCB: long-chain base
PA: phosphatidic acid
PI: phosphatidylinositol
PI3P: PI 3-phosphate
SL: sphingolipid
SNAP: soluble NSF adaptor protein
SNARE: soluble N-ethylmaleimide-sensitive factor attachment protein receptor
VLCFA: Very long-chain fatty acid
YPD: yeast extract, peptone, dextrose

## Current Addresses

Logan R. Hurst – The EVERY Company, Daly City CA, 94014

David A. Rivera-Kohr – Department of Biochemistry, University of Wisconsin, Madison WI, 53706

Adam Balutowski – Department of Biochemistry, Washington University, St. Louis MO 63110

Michael R. Hrabak – Department of Cellular & Molecular Pharmacology, Rosalind Franklyn University, Chicago IL 61801

Thomas D.D. Kazmirchuk – Department of Biology, Carlton University, Ottawa Ontario Canada

## Synopsis

Sphingolipids are known to stimulate the formation of rigid membrane domains that can affect protein function, however their role in vacuole fusion is not well understood. This study shows that sphingolipids containing 26-carbon very length fatty acids (VLCFAs) promote robust fusion by regulating the localization of the late endosomal Rab GTPase Ypt7 at the site of SNARE-mediated fusion.

- Elo3 adds the final two carbons to long chain fatty acids to make C-26 VLCFAs that are added to sphingolipids.
- Deletion of *ELO3* leads to abnormal vacuole morphology and an increase in membrane fluidity which are associated with the inhibition of vacuole fusion.
- The defect in *elo3*Δ vacuole fusion mapped to the tethering stage as shown by their inability to from large vesicles clusters.
- Altered tethering of *elo3*Δ vacuoles was due to the faulty distribution of the Rab GTPase Ypt7 which drives tethering through its interactions with the HOPS complex.

